# RNase L regulates the antiviral proteome by accelerating mRNA decay, inhibiting nuclear mRNA export, and repressing RNAPII-mediated transcription

**DOI:** 10.1101/2025.07.18.665572

**Authors:** J. Monty Watkins, Cameron J. Douglas, Renee Cusic, Ciaran P. Seath, James M. Burke

## Abstract

Ribonuclease L (RNase L) is an antiviral endoribonuclease that triggers widespread degradation of cellular mRNAs. Here, we show that the degradation of cellular mRNA by RNase L is a conserved response to flaviviruses, including Zika virus (ZIKV), dengue virus serotype 2 (DENV-2), and West Nile virus (WNV). Quantitative mass spectrometry in response to dsRNA or ZIKV infection shows that RNase L broadly downregulates the cellular proteome, reducing proteins with short half-lives involved in cell cycle progression, cellular metabolism, and protein synthesis. The mRNAs encoded by interferon-stimulated genes (ISGs) evade mRNA decay by RNase L, allowing for protein synthesis of ISG-encoding mRNAs. However, RNase L dampens ISG protein synthesis by triggering a block in nuclear mRNA export and repressing RNAPII-mediated transcription at later times during the antiviral response. These findings implicate reprograming of the cellular proteome as primary means by which RNase L combats viral infection, tumorigenesis, and immune dysregulation.

**HIGHLIGHTS:**

- RNase L initiates widespread decay of cellular mRNA in response to flaviviruses
- RNase L-mediated mRNA decay broadly downregulates the homoeostatic proteome
- Antiviral mRNAs are translated due to their ability to evade RNase L-mediated decay
- RNase L reduces antiviral proteins by inhibiting mRNA export and RNAPII-mediated transcription

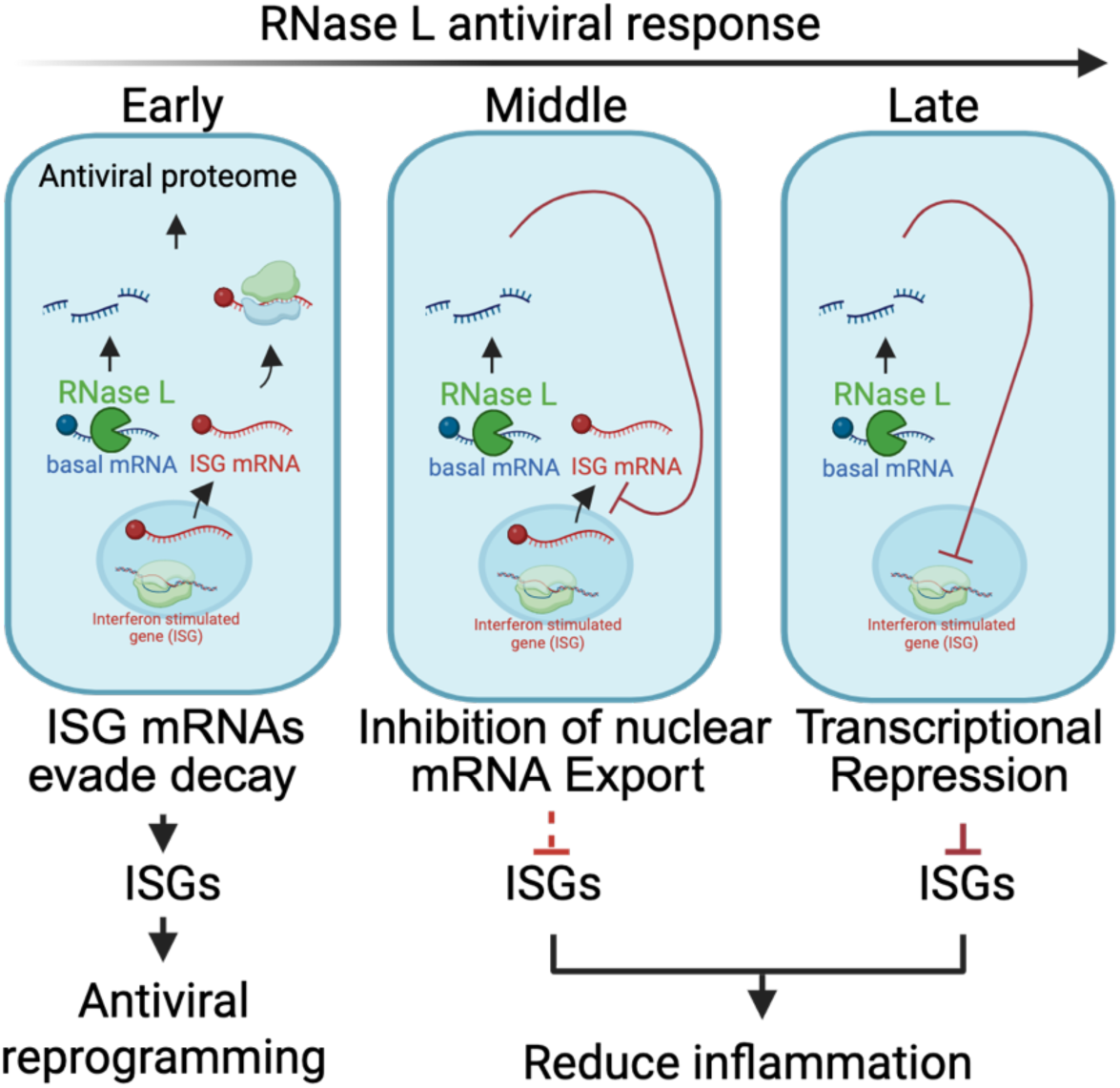

## INTRODUCTION

Ribonuclease L (RNase L) is a latent endoribonuclease constitutively expressed in most human cell types^1^. RNase L activates upon condensation of oligoadenylate synthetase 3 (OAS3) on viral dsRNA, which initiates the synthesis of pppA2’p5’A2’p5’A (2-5A). ^2–5^ 2-5A binds to the N-terminal ankyrin repeat domain (ARD) of RNase L, which allows for complementation of its C-terminal RNase domain via homodimerization.^6^ RNase L cleaves ssRNA at UN^N motifs in viral and host RNAs,^7,8^ which antagonizes the replication of numerous and diverse viruses.^9–14^ RNase L has been widely thought to suppress viral replication by cleaving viral RNA, cleaving ribosomes to arrests translation, and/or generating RNA fragments that induce the type I interferon response, autophagy, and the inflammasome.^15–18^

More recently, RNase L was shown to trigger widespread degradation of cellular mRNA.^19–21^ The loss of cellular mRNA was shown to be the primary means by which RNase L reduces bulk translation, as RNase L-cleaved ribosomes were shown to be translationally competent.^20^ Importantly, this allows for the translation of mRNAs that encode for proteins critical for the antiviral response, such as type I interferons (i.e., interferon beta; IFNB1), which evade RNase L-mediated mRNA decay.^19,22–24^ In addition, the rapid degradation of cytosolic mRNA by RNase causes RNA-binding proteins (RBPs) to translocate from the cytosol to the nucleus, which coincides with defects in nuclear RNA processing and inhibition of nuclear-cytoplasmic mRNA export.^25^ The inhibition of nuclear mRNA export occurs at later times post-activation of RNase L and reduces type I and type III interferon production.^23^ How RNase L-mediated mRNA decay and its secondary effects broadly alter the proteome during the antiviral response has not been investigated.

In this article, we characterize how RNase L alters the cellular proteome by quantitative mass spectrometry following dsRNA lipofection. These analyses show that RNase L broadly downregulates the homeostatic cellular proteome over the course of the innate antiviral response by accelerating mRNA decay, inhibiting export, and repressing RNAPII-mediated transcription. The mRNAs encoded by ISGs are induced to high levels during the initial phase of the RNase L response due to their ability to evade RNase L-mediated mRNA decay, thus allowing for the synthesis of these key antiviral proteins. However, RNase L-mediated inhibition of nuclear mRNA export and repression of RNAPII dampen protein production of ISGs over the course of the antiviral response. These findings establish how RNase L alters the cellular proteome to regulate cellular processes and inflammation in response to viral infection.

## RESULTS

### RNase L degrades mRNAs during viral infections and has minimal impact on apoptosis

The activation of RNase L in response to lipofection of poly(I:C), a synthetic viral dsRNA mimic, initiates widespread decay of cellular mRNA.^19,20^ Whether this occurs in response to viral infection, and what the frequency of this response is in individual cells, has not been thoroughly characterized. Therefore, we infected parental (WT) or RNase L-KO (RL-KO) A549 cells with dengue virus serotype 2 (DENV-2), Zika virus (ZIKV), or West Nile virus (WNV) and assessed human GAPDH mRNA and total poly(A)+ RNA cellular mRNA levels by single-molecule RNA in situ hybridization (smRNA-FISH). We also co-stained for viral RNA.

We observed an RNase L-dependent reduction in cytosolic *GAPDH* mRNA levels (∼100-fold) in the majority of A549-WT cells (66%-81%) infected with DENV-2, ZIKV or WNV, similar to poly(I:C) lipofection (**Figure 1A,B**). The loss of *GAPDH* mRNA corresponded to a reduction in cytosolic poly(A)+ RNA levels (**Figure 1A,C**). These data indicate that flaviviruses frequently trigger RNase L-mediated decay of cellular mRNA. Remarkably, viral RNA was relatively abundant in cells that activated RNase L-mediated mRNA decay, though RNase L generally reduced viral RNA loads (**Figure 1A,D**). This suggests that RNase L preferentially targets cellular mRNA and/or that viral RNAs evade RNase L-mediated RNA decay.

**Figure 1.**
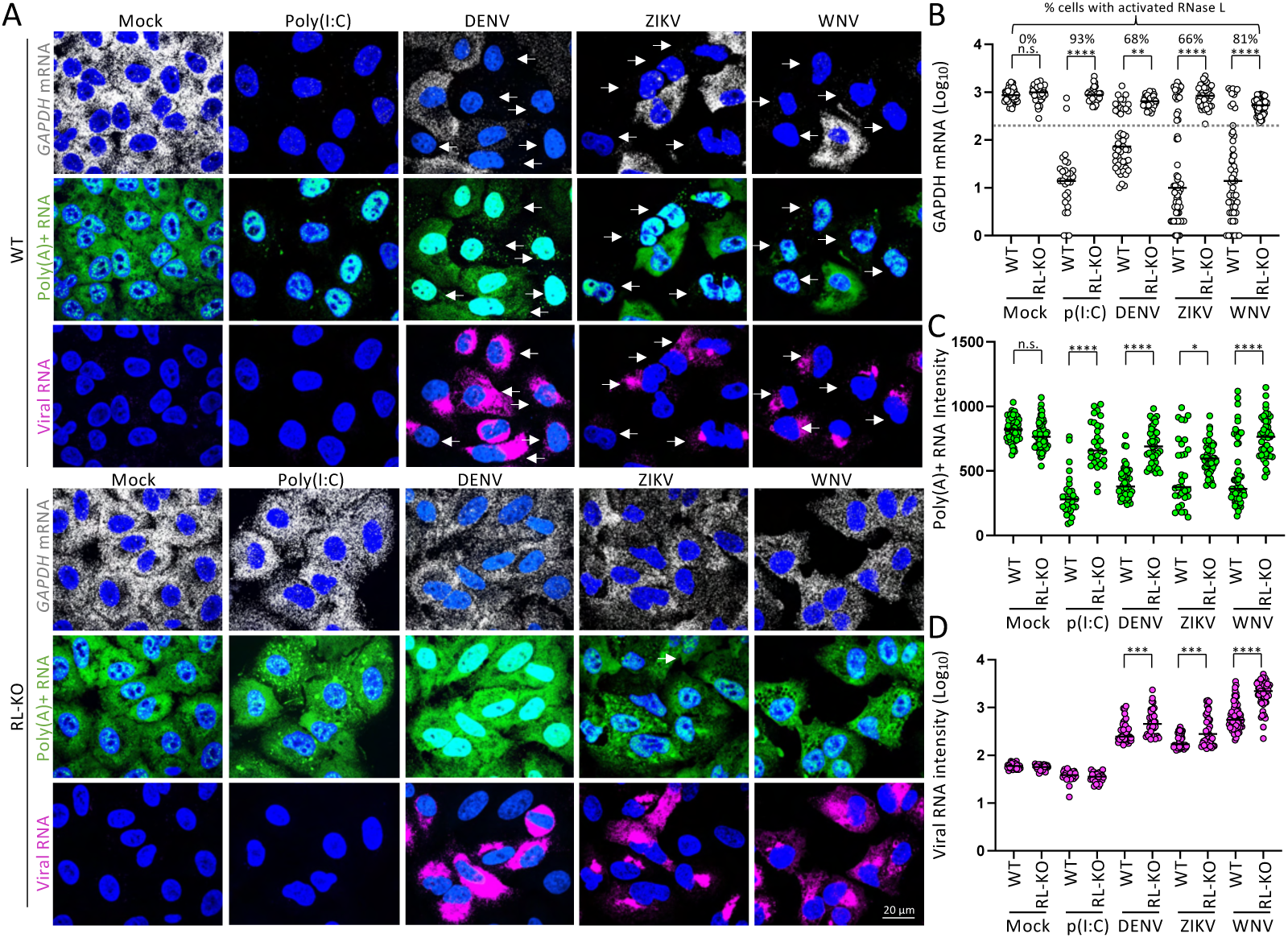
Flaviviruses trigger RNase L-mediated decay of cellular mRNA. (A) smRNA-FISH for human GAPDH mRNA (gray) and viral RNA (magenta) in parental (WT) or RNase L-KO (RL-KO) A549 cells 24 hours post-infection (p.i.) with DENV-2, ZIKV, or WNV. As a control, cells were lipofected with poly(I:C) for 12 hours. Arrows indicate cells that activate RNase L-dependent decay of cellular mRNA. Poly(A)+ RNA staining for these panels is shown in Fig. S1A. (B) quantification of GAPDH mRNA as represented in (A) in individual cells (dot). The percentage of cells with activated RNase L was determined by the percent of cells with GAPDH mRNA copies below the lowest observed in RL-KO cells (dashed line). The bar represents the median. (C) Quantification of poly(A) RNA intensity as represented in (A). (D) Quantification of viral RNA copies as represented in (A).

### Cells remain viable following RNase L-mediated decay of cellular mRNA

RNase L is thought to initiate apoptosis.^15,26,27^ To examine how long cells remain viable after initiating RNase L-mediated mRNA decay, we performed live-cell imaging of poly(A)-binding protein C1 (PABPC1) tagged with mRuby2 (mRuby2-PABPC1), which assembles into RNase L-induced bodies (RLBs) and translocates to the nucleus as RNase L initiates the degradation of cellular mRNA^28^. As expected, PABPC1 assembled into RLBs and translocated from the cytosol to the nucleus following lipofection of poly(I:C) in WT cells but not RNase L-KO cells, which formed PABPC1-positive stress granules as the result of protein kinase R (PKR) activation^28^ (**Figure 2A**).

**Figure 2.**
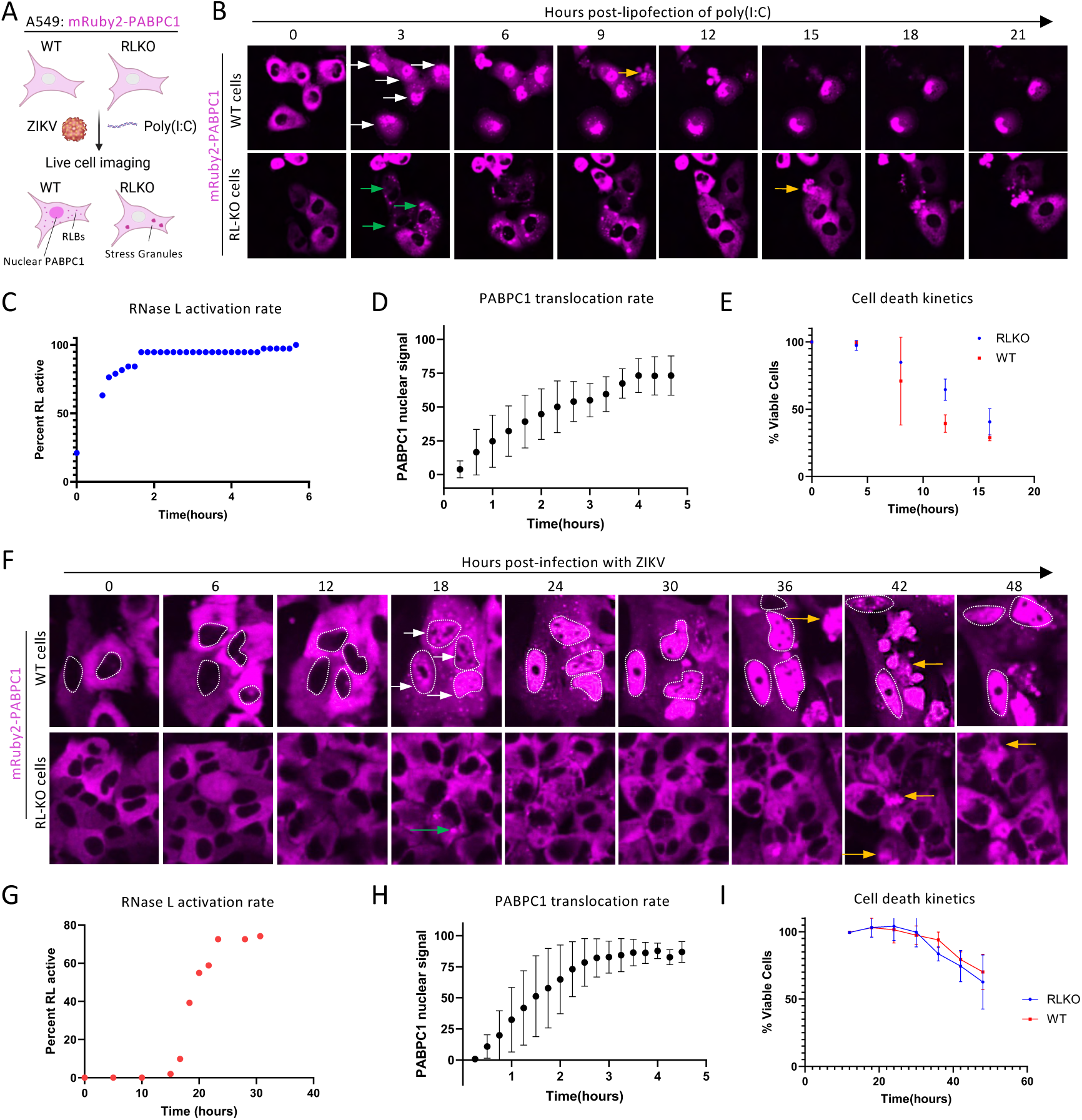
RNase L kinetics in response to poly(I:C) and ZIKV infection. (A) Schematic of experimental design. (B) Live cell imaging of mRUBY2-PABPC1 A549 cells during poly(I:C) transfection. White arrows indicate nuclear PABPC1, green arrows indicate stress granules, and orange arrows indicate dead cells. (C) Percentage of cells with RNase L active in a single field of view, as determined by initiation of PABPC1 translocation. (D) Intensity of nuclear PABPC1, normalized to the peak value in each individual cell. Error bars represent standard deviation for n = 10 cells. (E) Kinetics of cell viability in response to poly(I:C) from 2 biological replicates. (F) Live cell imaging of mRUBY2-PABPC1 A549 cells during ZIKV infection. White arrows indicate nuclear PABPC1, green arrows indicate stress granules, and orange arrows indicate dead cells. (G) Percentage of cells with RNase L active in a single field of view, as determined by initiation of PABPC1 translocation (H) Intensity of nuclear PABPC1, normalized to the peak value in each individual cell. Error bars represent standard deviation for n = 10 cells. (I) Kinetics of cell viability in response to ZIKV from 2 biological replicates.

We observed that most WT cells (>90%) initiated RNase L-dependent PABPC1 translocation by 2 hours post-lipofection of poly(I:C) (**Figure 2B,C**), with these cells completing PABPC1 translocation by 4 hours (**Figure 2B,D**). Thus, RNase L completes degradation of cellular mRNA within two hours post-activation. Remarkably, many of these cells lived for several hours after completing RNase L-mediated decay of cellular mRNA (**Fig. 2B,E**). Moreover, WT and RL-KO cells died at a comparable rate in response to poly(I:C) lipofection (**Fig. 2B,E**), indicating that RNase L does not accelerate cell death in response to dsRNA lipofection.

We performed similar analyses following Zika virus infection. RNase L-dependent translocation of PABPC1 from the cytosol to the nucleus initiated between 18- and 24-hours p.i. (**Figure 2F,G**), and the rate of PABPC1 translocation to the nucleus following RNase L activation, as determined by RLB assembly, in response to ZIKV was similar to that observed in response to poly(I:C), whereby PABPC1 completed translocation within ∼2 hours (**Figure 2H**). These, data indicate that RNase L activates in individual ZIKV-infected cells between 18- and 24-hours p.i. and degrades nearly all cellular RNA within 2 hrs. following activation. Nevertheless, ZIKV-infected cells remained viable for several hours following RNase L activation, with some cells remaining viable over 24 hours (**Figure F,I**). Moreover, we did not observe a difference cell death kinetics between WT and RL-KO cells over the course of ZIKV infection (**Figure 2F,H** and **Figure S1**). These data demonstrate that A549 cells remain viable following RNase L-mediated decay of cellular mRNA during ZIKV infection.

### RNase L broadly alters the cellular proteome during the innate immune response

Because most cells remain viable following RNase L-mediated mRNA decay, we sought to examine how RNase L-mediated mRNA decay broadly remodels the over the course of the innate immune response. To do this, we performed quantitative mass spectrometry in WT and RL-KO A549 cells under mock conditions or following poly(I:C) lipofection (8 or 16 hours). We did not observe substantial differences between the cellular proteome in WT and RL-KO cells under mock conditions (**Figure S2A**), and we only observed minor alterations to the proteome at 8 hrs. post-lipofection of poly(I:C) (**Figure S2B**).

At 16 hours post-lipofection of poly(I:C), we observed substantial differences in the proteome between WT and RL-KO cells (**Figure 3A**). We conducted unpaired t-tests of WT and RL-KO cells to identify proteins that were significantly up- or down-regulated in an RNase L-dependent manner at 16 hours post-lipofection of poly(I:C) (**Figure 3B** and **Data S1**). As expected, ISG proteins (i.e., ISG15, IFIT1, MX1, DDX58) were significantly upregulated in both WT and RL-KO cells following poly(I:C) lipofection. We observed 447 proteins significantly downregulated in an RNase L-dependent manner, and 274 proteins upregulated in an RNase L-dependent manner (**Figure 3C** and **Data S1**).

**Figure 3.**
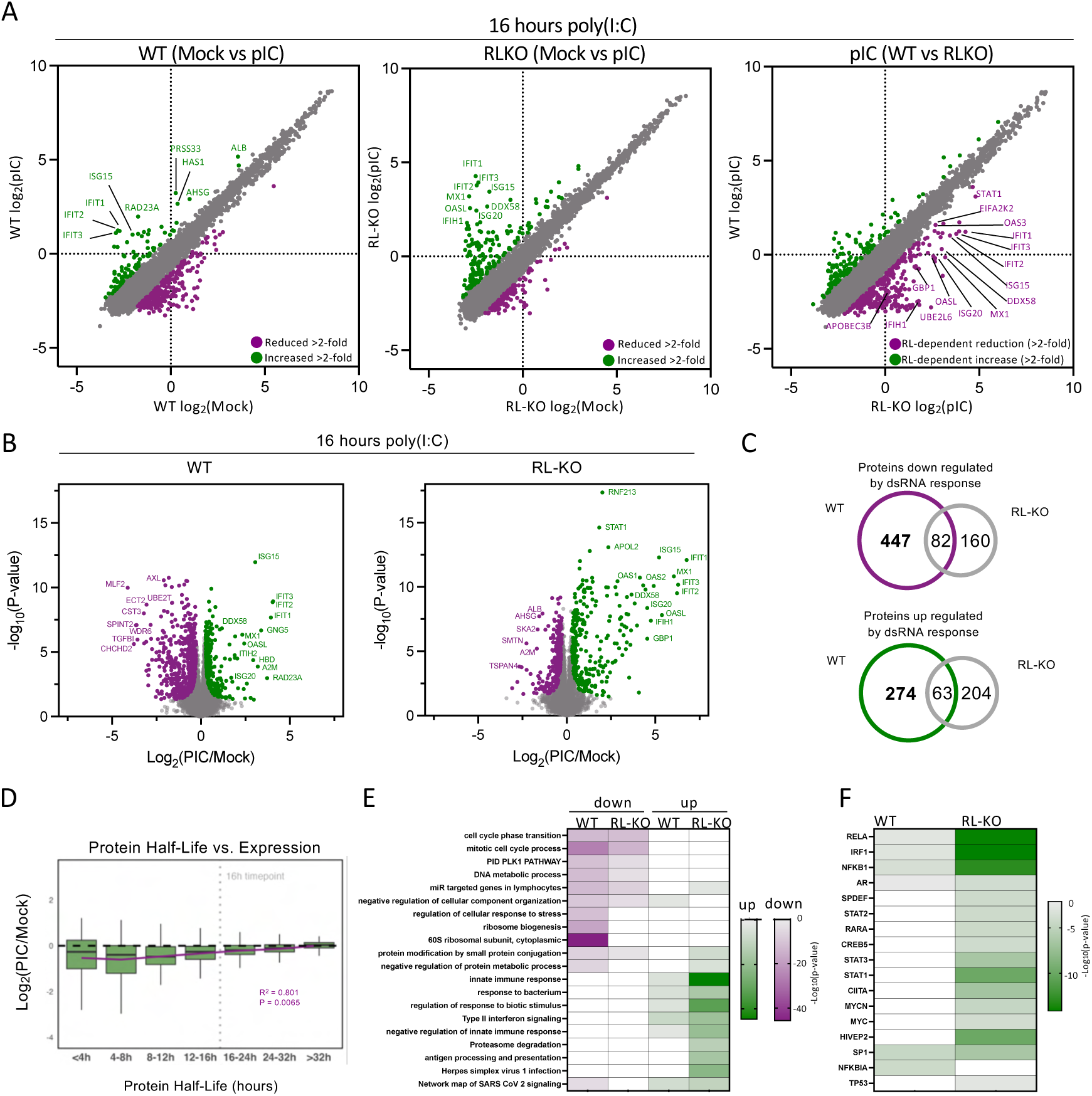
Broad effects of RNase L on the cellular proteome. (A) Scatter plot protein levels in mock conditions (x-axis) and 16 hours post-lipofection of poly(I:C) in WT or RL-KO cells. (B) Volcano plots showing proteins up- or down-regulated in WT or RL-KO cells at 16 hours post-lipofection of poly(I:C). (C) Venn diagram of proteins that are significantly differentially regulated by dsRNA response. Significance determined by two sample T-test with FDR<0.05 (n=6). (D) Box-plot of the Log2FC of proteins within indicated half-life cutoff. Line is linear regression of mean Log2FC for each half-life cutoff with R^2^=0.801 and P= 0.0065. (E) Heatmap of GO terms associated with the proteins that were significantly upregulated or downregulated in indicated cell line 16 hours post-lipofection of poly(I:C). (F) Heatmap of transcriptional regulatory relationships of differentially upregulated proteins in WT and RL-KO cells 16 hours post-lipofection of poly(I:C).

We considered the possibility that the proteins downregulated by RNase L is due to them having short half-lives, thus leading to their more rapid reduction following the degradation of cellular mRNAs by RNase L. Indeed, cross-referencing the downregulated proteins with a published database of protein half-lives showed that proteins with short half-lives were downregulated by RNase L in comparison to proteins with long half-lives, which were largely unaffected by RNase L (**Figure 3D**). These data suggest that loss of protein synthesis plays a significant role in the function of RNase L, whereby the arrest of protein synthesis via mRNA decay broadly downregulates the cellular proteome, which is correlated with the inherent stability of specific proteins rather than the activation of a specific protein degradation mechanisms.

We performed gene ontology (GO) analyses on proteins up- or down-regulated in WT and RL-KO cells (**Figure 3E**). Both cell lines downregulated proteins involved in cell cycle and upregulated proteins involved in innate immunity to viral infection. Notable processes enriched among RNase L-downregulated proteins involved regulation of cellular stress and ribosome biogenesis. A particularly important finding was that RNase L selectively regulated transcriptional activity by TRRUST analysis. The activity of several transcription factors, including STAT1, MYC, AR, and CREB5 was downregulated by RNase L, whereas proteins controlled by NFKB1, NFKBIA, SP1, IRF1, and RELA were upregulated in both WT and RL-KO cells (**Figure 3F**).

### RNase L dampens the production of proteins encoded by interferon-stimulated genes

Although the activation of RNase L results the degradation of nearly all cellular mRNAs, the mRNAs induced by immediate early genes have been shown to [i.e., interferon beta (IFNB1)] accumulate to abundant levels, which critical for the synthesis of key antiviral proteins.^19,20^ However, it is currently unknown whether the mRNAs encoded by interferon-stimulated genes (ISGs) similarly evade RNase L-mediated mRNA decay and how RNase L broadly alters ISG protein synthesis.

#### RNase L downregulates ISG proteins

To investigate how RNase L-mediated mRNA decay affects in induction of ISG proteins, we plotted the differential (Mock vs pI:C) between WT and RL-KO cells at 8 and 16 hours post-lipofection of poly(I:C) (**Figure 4A**). We observed an increase (2- to 32-fold) in a modest number of proteins involved in antiviral defense in both WT and/or RL-KO cells, such as GBP1, IFIT3, IFIT1, ISG15, OASL, IRF7, IFNL1, and IFNL3 (**Figure 4A,B)**. At 16 hours post-lipofection of poly(I:C), both WT and RL-KO cells induced many antiviral proteins, including IFITs, MX1, ISG15, OASL, GBP1, ISG20, OAS2, IFIH1 (MDA-5), and DDX58 (Rig-I) (**Figure 4A,B**).

**Figure 4.**
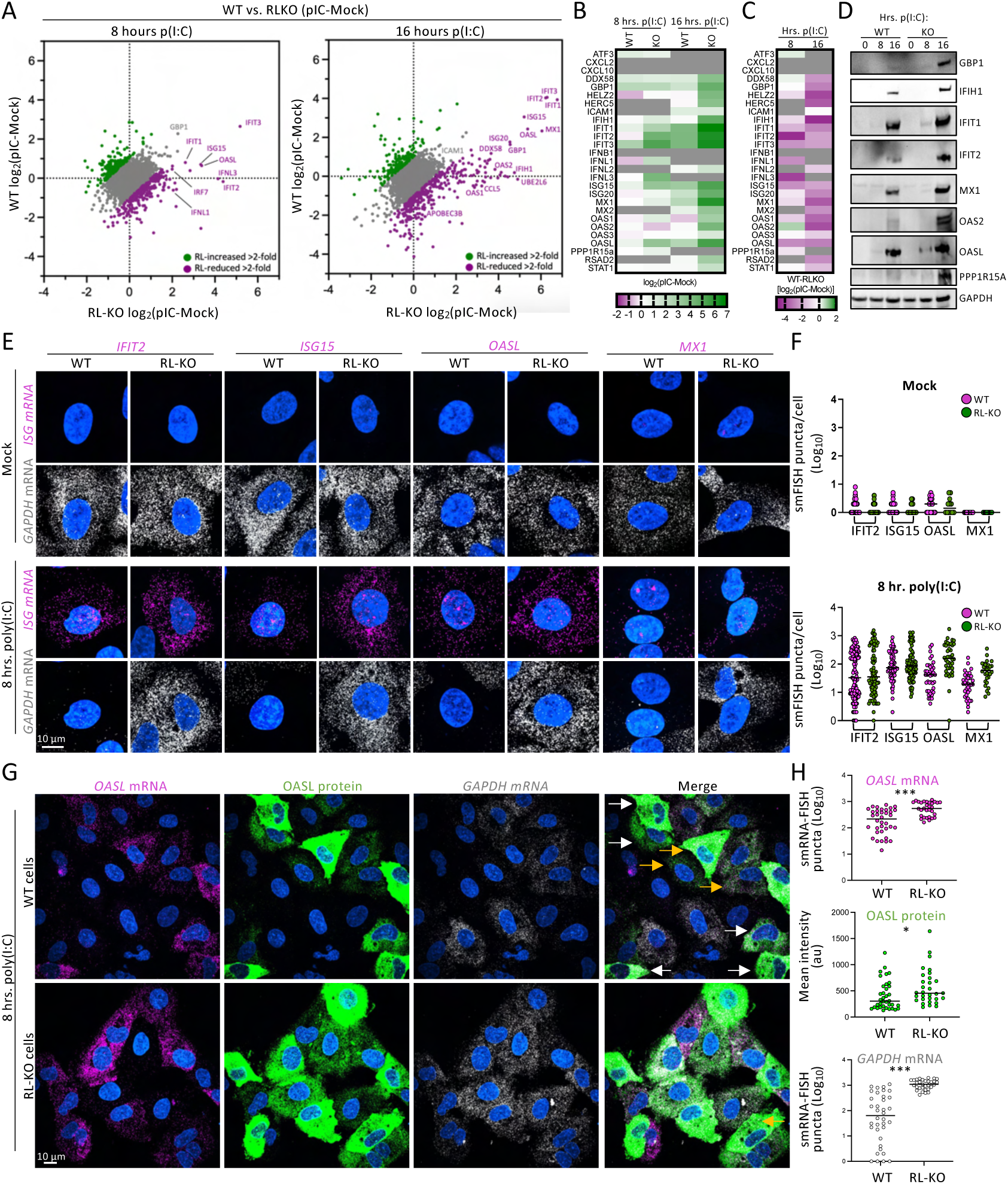
ISG mRNAs evade RNase L-mediated RNA decay, permitting their translation. (A) Quantitative proteomic analyses in WT and RL-KO cells. (B) Heat map showing the induction of immediate early and ISGs in response to poly(I:C). (C) Heat map showing the effect of RNase L from (B). (D) Western blot analysis of ISG proteins in WT and RL-KO at 0, 8, and 16 hours post-lipofection of poly(I:C). (E) smRNA-FISH for indicated ISG mRNAs and GAPDH mRNA under mock conditions or 8 hours post-lipofection of poly(I:C). White arrows indicate cells with degraded *GAPDH* mRNA, yellow arrows indicate cells with *GAPDH* mRNA. (F) Quantification of smRNA-FISH puncta as represented in (E). Each dot represents a single cell. (G) smRNA-FISH for *OASL* mRNA and *GAPDH* mRNAs and IFA for OASL protein. (H) Quantification of smRNA-FISH puncta and mean fluorescence intensity (MFI) for OASL protein in individual cells (dots) as represented in (G).

Importantly, nearly all antiviral proteins were higher in RL-KO cells in comparison to WT cells at both 8 and 16 hours post-lipofection of poly(I:C), though the difference was more pronounced at 16 hours (**Figure 4C** and **Figure S3A,B**). Western blot analyses corroborated our MS data, showing that GBP1, IFIH1, IFIT1, IFIT2, MX1, OAS2, OASL, and PPP1R15A (GADD34) increased in both WT and RL-KO cells following lipofection of poly(I:C), yet they were all lower in WT cells in comparison to RL-KO cells (**Figure 4D**). Notably, we did not readily detect some antiviral proteins (GBP1, IFITs, MX1, OAS2) at 8 hours post-lipofection of poly(I:C) in either WT or RL-KO cells (**Figure 4D**), consistent with their lower levels and/or lack of detection in our MS analyses.

#### ISG-encoding mRNAs evade RNase L-mediated mRNA decay and are translated

Over 90% of A549 cells activate RNase L-mediated mRNA decay by 6 hours post-lipofection of poly(I:C) (**Figure 2B,C**). However, based on our analyses, as well as previous analyses of type I/III interferon secretion,^23^ antiviral protein synthesis occurs between 8 and 16 hours. Therefore, we wanted to determine if the mRNAs encoding ISGs evade RNase L-mediated mRNA decay. To assess this, we lipofected WT or RL-KO cells with poly(I:C) for 8 hours and performed smRNA-FISH for the mRNAs encoding *IFIT2*, *ISG15*, *OASL*, or *MX1* and *GAPDH* mRNA.

In mock-treated WT and RL-KO cells, the mRNAs encoding *IFIT2*, *ISG15*, *OASL*, or *MX1* were lowly expressed (<10 copies), with most cells containing no detectable mRNAs (**Figure 4E**). However, these mRNAs were induced in cells following lipofection of poly(I:C), with individual cells expressing between 10-1000 copies. Importantly, WT cells that lacked *GAPDH* mRNA contained abundant levels of these ISG-encoding mRNAs. Average IFIT2 and ISG15 mRNA levels were comparable between WT and RL-KO cells (**Figure 4E,F**), whereas MX1 and OASL mRNAs were lower on average n WT cells in comparison to RL-KO cells. These data indicate that the mRNAs encoding ISGs can evade RNase L-mediated mRNA decay, albeit to a varying degree.

To confirm that the protein synthesis of ISGs occurs in cells with activated RNase L, we performed smRNA-FISH for *OASL* and *GAPDH* mRNAs and co-stained for OASL protein by IFA. Consistent with our MS and western blot analyses, we did not observe OASL protein in cells under mock conditions (**Figure S4A**). However, at 8 hours post-lipofection of poly(I:C), we observed high OASL protein levels in many WT cells (**Figure 4G**). Importantly, most of these cells contained high levels of *OASL* mRNA while lacking *GAPDH* mRNA (**Figure 4G** and **Figure S3A**). On average, OASL protein levels were only slightly lower (∼twofold) in WT cells in comparison to RL-KO cells (**Figure. 4H**), which was comparable to the twofold lower *OASL* mRNA levels in WT cells. Notably, *OASL* mRNA and protein levels were positively correlated in both WT and RL-KO cells (**Figure S4B**). Similar results were obtained for IFIT2 (**Figure S4D**). These data demonstrate ISG mRNAs effectively evade RNase L-mediated mRNA decay, which is sufficient for their translation.

#### RNase L-dependent nuclear retention of ISG mRNAs dampens ISG protein synthesis

We next addressed how RNase L limits ISG protein production at late times (16 hrs.) post-lipofection of poly(I:C). Previous studies showed that RNase L triggers a block in canonical nuclear export of *IFNL1* mRNA between 8 and 16 hours post-lipofection of poly(I:C), resulting in a reduction in IFNL1 protein synthesis. Therefore, we examined if ISG proteins are reduced due to the retention of their mRNAs in the nucleus.

We observed that many ISG-encoding mRNAs (OASL, IFIT2) are retained in the nucleus at 16 hours post-lipofection of poly(I:C), whereby their nuclear levels increased on average 5-fold in comparison to RL-KO cells while their cytosolic levels decreased (**Figure 5A,B**). However, some ISG mRNAs, such as *ISG15*, were not frequently or robustly retained in the nucleus (**Figure 5A,B**). Consistent with this notion, WT cells co-stained for *ISG15* and *OASL* mRNAs displayed nuclear retention of *OASL* mRNA but not *ISG15* mRNA (**Figure S4E**). These data suggest that RNase L differentially affects mRNA export, as previously observed for *IFNL1* and *IFNB1* mRNAs.^23^

**Figure 5.**
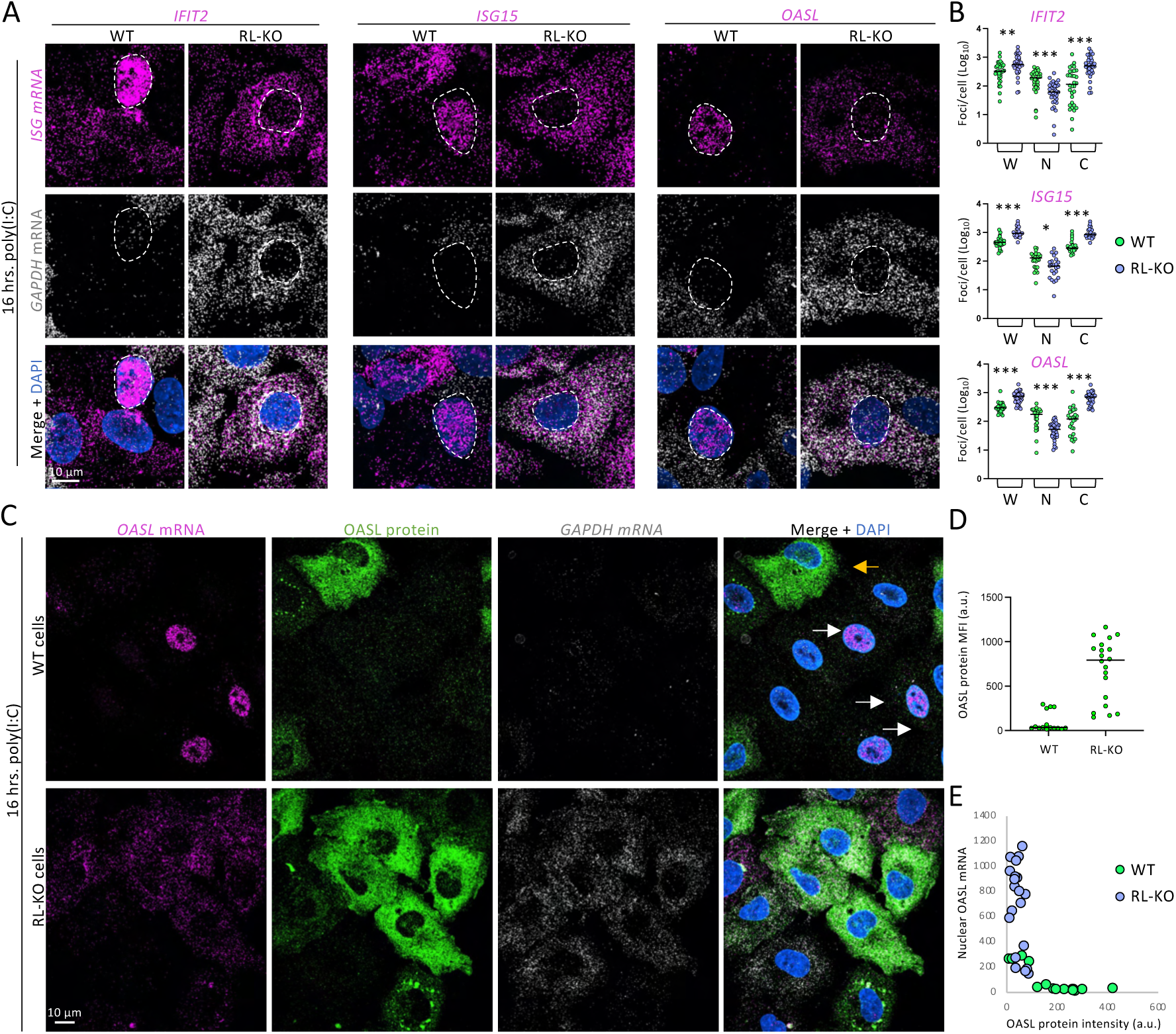
RNase L-dependent inhibition of ISG mRNA export dampens protein synthesis. (A) smRNA-FISH for *IFIT2*, *ISG15*, or *OASL*, and *GAPDH* mRNAs 16 hours post-lipofection of poly(I:C) in WT and RL-KO cells. The dashed line demarcates nuclei, as determined by DAPI staining (blue), of cells displaying a retention of mRNAs. (B) Quantification of total (W), cytosolic (C), or nuclear (N) ISG mRNA levels as represented in (A). (C) Co-IFA for OASL protein and smRNA-FISH for OASL mRNA. (D) Mean fluorescence intensity (MFI) of OASL protein in WT and RL-KO cells as represented in (C). (E) Scatter plot of the mean fluorescence intensity (MFI) of OASL protein (x-axis) and nuclear OASL mRNA (y-axis) per cell as represented in (C).

To determine if the nuclear mRNA export reduced protein levels at 16 hours post-lipofection of poly(I:C), we performed co-IFA for *OASL* mRNA and IFA for OASL protein (**Figure 5C**). These analyses demonstrated that WT cells contained significantly lower OASL protein than RL-KO cells (**Figure 5C,D**), and that the cells displaying nuclear retention of *OASL* mRNA contained the lowest OASL protein levels (**Figure 5C,F**). These data support that RNase L-mediated inhibition of nuclear export dampens the protein synthesis of ISGs at later times post-poly(I:C).

#### RNase L activation leads to the repression of RNAPII-mediated transcription

Our data show that total ISG mRNA levels are reduced at late times (16 hrs.) post-lipofection of poly(I:C) in an RNase L-dependent manner, due in part to accelerated mRNA decay and nuclear mRNA export inhibition. However, we considered the possibility that RNase L could trigger a reduction in RNAPII-mediated transcription. The rationale for this hypothesis is that the translocation of RNA-binding proteins from the cytosol to the nucleus in response to MHV-68 SOX (muSOX)-mediated mRNA decay inhibits RNAPII-mediated transcription.^29^ Moreover, we observed a reduction in ISG15 protein synthesis despite the lack of ISG15 mRNA retention in the nucleus. Therefore, we examined if RNase L activation reduces RNAPII-mediated transcription and whether this would affect the transcriptional induction of immediate early antiviral genes or ISGs such as ISG15.

To test whether RNase L activation could repress RNAPII transcription, we measured transcriptional activity in WT and RL-KO cells by pulsing them with 5-Ethynyl Uridine (EU). We quantified EU incorporation by fluorescence microscopy using Alexa Fluor-488 (EU-click-488). We also performed IFA for poly(A) binding protein C1 (PABPC1) as a marker for RNase L activation, as PABPC1 translocates to the nucleus upon RNase L activation. In both WT and RL-KO cells, incorporation of EU (EU+) resulted in intense nuclear fluorescence upon click chemistry of Alexa Fluor-488 (EU-click-488) in comparison to cells lacking EU (EU-), and treatment with actinomycin D reduced EU-click-488 staining (**Figure 6A,B** and **Figure S5A**). These data demonstrate that EU-click-488 fluorescence is dependent on RNAPII-mediated transcription.

**Figure 6.**
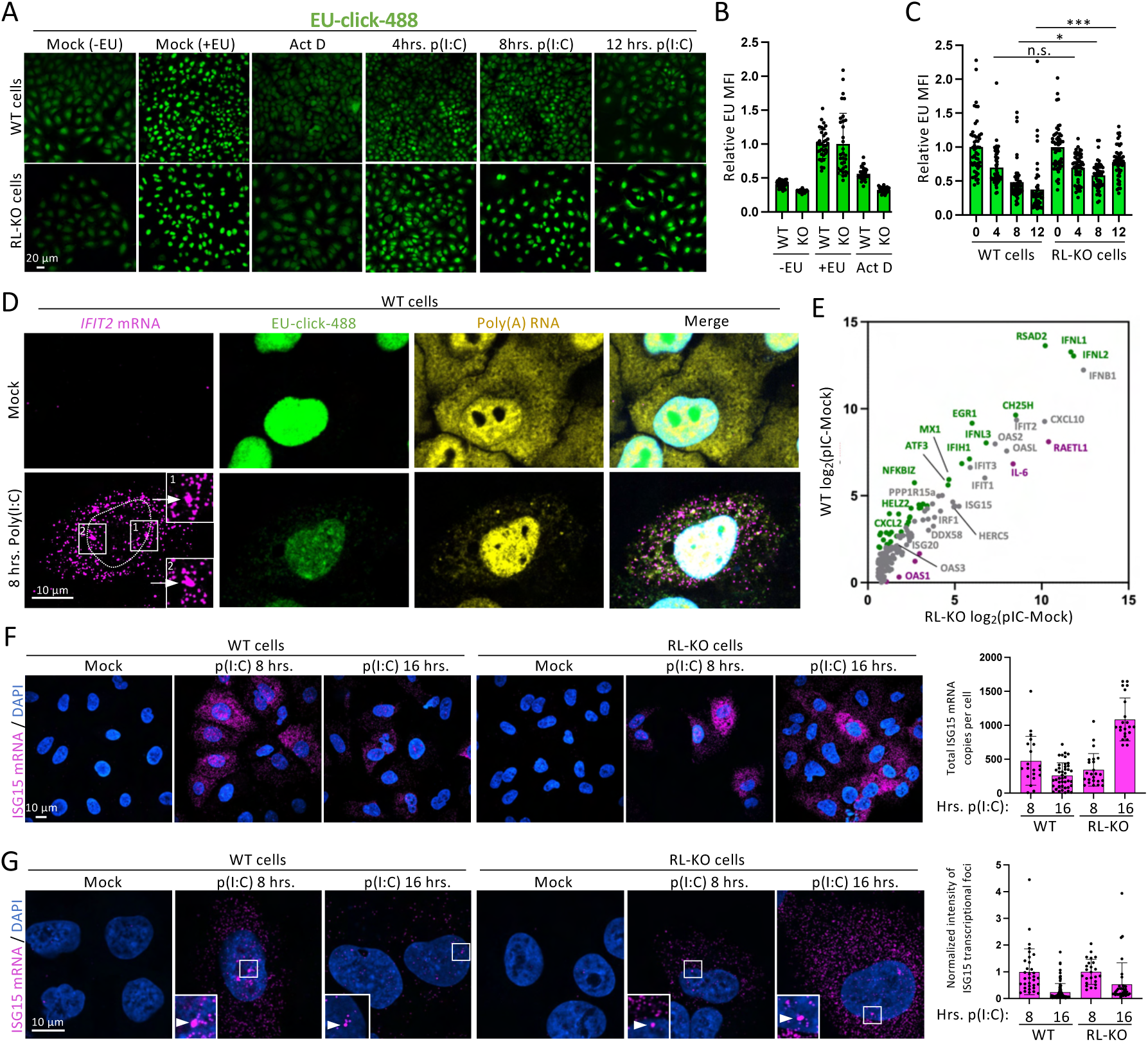
RNase L dampens ISG expression by repressing RNAPII-mediated transcription. (A) Fluorescence microscopy of EU-click-488 in WT and RL-KO cells. (B and C) Quantification of EU mean fluorescence intensity (MFI) in cells (dots). The bars represent the average +/- standard deviation. (D) smRNA-FISH for IFIT2 mRNA and poly(A)+RNA and in WT cells containing EU-click-488 under mock treatment or 8 hours post-lipofection of poly(I:C). The arrow in the inset indicates putative nascent mRNAs at the genomic locus. (E) Normalized EU intensity in cells with or without stress granules. (F) Meta analyses of RNA-seq data of RNA induced in WT and RL-KO A549 cells 6 hours post-lipofection of poly(I:C) (Burke et al., 2019). The mRNAs upregulated (green), unchanged (gray), or downregulated (Magenta) RNase L are indicated. (F) smRNA-FISH for ISG15 mRNA in WT and RL-KO cells at 0-, 8-, or 16-hrs. post-lipofection of poly(I:C). The graph on the right is quantification of total ISG15 mRNA copies per cell. (G) Similar to (F) but displayed to show nuclear ISG15 transcription foci (arrow). The graph on the right is the integrated density of ISG15 transcription foci normalized to the 8 hrs. post-lipofection of poly(I:C) time point.

In WT cells, we observed a progressive reduction in EU staining following the lipofection of poly(I:C) (**Figure 6A,C**). At 4 and 8 hours, EU staining was reduced by 25% and 50%, respectively. By 12 hours, it was reduced by greater than 60%. This reduction was RNase L-dependent, as EU staining was significantly higher in RL-KO cells (**Figure 6A,C**). Moreover, EU staining in WT cells containing nuclear PABPC1 (RNase L active) was lower than in WT cells displaying cytosolic PABPC1 (RNase L inactive) (**Figure S5A,B**). These data support the notion that RNase L activation correlates with a reduction in RNAPII-mediated transcription, particularly at late times post dsRNA stress (12 hrs.). We note that RL-KO cells displayed a reduction (∼25%) in EU staining throughout the time course (**Figure. 6A,C**), suggesting that an additional innate immune response to dsRNA represses RNAPII-mediated transcription. Notably, these cells tended to contain PABPC1-positive stress granules (**Figure S5B,D**), suggesting that the PKR pathway, the integrated stress response (ISR), and/or stress granules also repress RNAPII-mediated transcription.

#### ISGs are induced despite repression of bulk RNAPII-mediated transcription at early times

Three observations indicate that antiviral genes – immediate early genes, such as type I interferons and interferon-stimulated genes (ISGs) – are induced despite global repression of RNAPII-mediated transcription mediated by RNase L. First, our data showed that cells with activated RNase L contain ISG-encoding mRNAs (**Figure 3E**). Second, *IFIT2* mRNA was abundant in cells that activated RNase L (reduced cytosolic poly(A)+ RNA) and displayed transcriptional repression (reduced EU) at 8 hours post-lipofection of poly(:C) (**Figure 6D, S6**). Importantly, we observed two nuclear structures consistent with nascent *IFIT2* RNA transcripts at the *IFIT2* genomic loci (**Figure 5D**),^19^ supporting that the *IFIT2* gene is actively transcribed despite bulk repression of transcription. Notably, the reduction of nuclear EU staining by RNase L is not due to the degradation of nuclear RNA, as poly(A)+ RNA was not reduced in the nucleus in cells that activated RNase L. This is consistent with previous studies showing that RNase L is localized to the cytosol and does not degrade nuclear RNA.^21,25,30^ Third, consistent with the notion that IFIT2 mRNA is transcribed despite repression of RNAPII transcription, meta-analyses of RNA-seq from WT and RL-KO six hours post-lipofection of poly(I:C) showed that mRNAs encoding antiviral proteins, including IFIT2 as well as IFNB1, IFNL1, RSAD2, and OASL, are induced to comparable levels in WT and RL-KO cells (**Figure 6E**).^19^ These data show that ISGs can be transcribed, at least initially, in cells that have initiated RNase L-dependent repression of global transcription.

#### RNase L-mediated repression of RNAPII dampens ISG mRNA production at late times

We considered the possibility that RNase L-dependent repression of RNAPII-mediated transcription could eventually shut off ISGs, and that this could contribute to the broad reduction in ISG protein synthesis. Two observations support this hypothesis. First, we observed a reduction in total ISG15 mRNA from 8 hours to 16 hours poly(I:C) in WT but not RL-KO cells (**Figure 6F**). Second, we assessed transcriptional output in cells by measuring the integrated density of nascent transcripts at the genomic locus of *ISG15*. Typically, cells stained for *ISG15* mRNA by smRNA-FISH contained two intense foci in the nucleus (**Figure 6G** and **Figure S7A**). These foci were typically not observed in mock treated cells (**Figure S7A**), consistent with these cells lacking ISG15 mRNA due to a lack of transcriptional induction, and their intensity reduced following actinomycin D treatment (**Figure S7B**). Thus, these data indicate that these foci represent nascent ISG15 transcription. We term these transcription foci. Importantly, the intensity of ISG15 transcription foci reduced from 8 to 16 hours. post-lipofection of poly(I:C) in WT cells (**Figure 6G**). The reduction in the transcription foci intensity was more pronounced in WT cells than in RL-KO cells, though RL-KO cells also displayed a modest yet noticeable reduction in transcription foci intensity, consistent with the observation that cells can also reduce transcription in and RNase L-independent manner in response to poly(I:C) (**Figure 6A,C**). Overall, these data support that RNAPII-mediated transcription of ISGs reduces at late times during the innate antiviral response and that RNase L promotes this downregulation.

## DISCUSSION

In this article, we investigated how RNase L alters the cellular proteome during the innate antiviral response. Combined with previous studies,^19,20,25^ our data support a model whereby RNase L activation results in the rapid decay of cellular mRNA, which eventually leads to a block in mRNA export and repression of RNAPII-mediated transcription. These processes serve as a molecular timer that shuts down cellular protein synthesis in a hierarchical manner, whereby translation of basally expressed mRNAs is immediately arrested while the mRNAs encoding antiviral proteins evade RNase L-mediated degradation are translated. However, subsequent inhibition of nuclear mRNA and repression of RNAPII-mediated transcription mediated by RNase L reduces antiviral protein synthesis, providing a crucial regulatory mechanism that can dampen damaging inflammation.

Our data solidify that RNase L-mediated degradation of cellular mRNA is a frequent response to viruses that trigger RNase L activation, such as flaviviruses (**Figure 1A-C**). By monitoring PABPC1 localization using live cell imaging, we observed that the activation of RNase L is delayed until 18-24 hours post-infection with ZIKV (**Figure 2F,G**). However, once activated, RNase L degrades cellular mRNA to completion within individual cells within ∼2 hours (**Figure 2H**), resulting in most cells lacking basally expressed mRNAs by 24 hours p.i.. Remarkably, most of these cells survive for an additional 24 hours, and RNase L-KO cells died at a comparable rate as cells that activated RNase L in response to ZIKV (**Figure 2F,I**). Combined, these data demonstrate that neither the activation of RNase L nor degradation of cellular mRNA is immediately cytotoxic to cells. Because these observations contrast with many studies proposing that RNase L initiates apoptosis signaling cascades,^24,27^ we are actively investigating stress response and apoptotic pathways initiated by RNase L activation in biologically relevant contexts.

Our proteomic analyses demonstrate that RNase L-mediated mRNA decay leads to the broad downregulation of the cellular proteome (**Figure 3A-C**). Importantly, these data revelated that RNase L also represses antiviral protein synthesis, particularly at late times post-lipofection of poly(I:C) (**Figure A-H**). This is an important finding because RNase L is typically thought to promote the type I interferon response by generating cleavage fragments that activate Rig-I-mediated MAVS signaling,^16^ though subsequent studies demonstrated that cells with intact or robust RNase L responses dampen the protein synthesis of type I interferons.^31^ Combined with our previous studies and data herein, we propose that RNase L shuts down cellular processes in a stepwise manner that differentially alters the ability of cells to synthesize antiviral proteins. During the early phase of the RNase L response, ISG-encoded mRNAs can be transcribed, exported to the cytosol, and translated, as demonstrated for OASL and IFIT2 (**Figure 4E-H** and **Figure S4D**). Thus, similar to mRNAs encoded by immediate early genes (interferon beta),^19^ ISG-encoding mRNAs also evade RNase L-mediated mRNA decay, in effect, as RNase L cannot greatly increase the decay rate of these inherently unstable mRNAs. During the mid-phase of the RNase L response, nuclear export of ISG mRNAs is inhibited in an RNase L dependent manner (**Figure 5A,B**), which represses ISG protein synthesis (**Figure 5C**). During the late phase of the RNase L response, RNase L represses RNAPII-mediated transcription (**Figure 6A-C**). While ISGs are initially transcribed despite this broad reduction in RNAPII-mediated transcription (**Figure 6D,E**), the transcription of ISGs reduces at late time post-lipofection of poly(I:C) (**Figure 6F,G**). Thus, if ISG mRNAs are not induced before RNase L arrests mRNA export and RNAPII-mediated transcription, their ability to be translated is nearly abolished (**Figure 5C,D**). These observations reveal that the activation kinetics of the RNase L response relative to the type I interferon response likely determines how strongly RNase L represses the ISG response. Moreover, this could potentially be context-dependent, as viruses differentially trigger and/or antagonize these pathways. Future work will focus on understanding the relative kinetics of these pathways in response to diverse viruses and how this impacts the ISG response.

Our data herein support a novel mechanism by which RNase L dampens host gene expression, which is the repression of RNAPII-mediated transcription **(Figure 6A-C**). The ability of RNase L-mediated mRNA decay in the cytosol to regulate nuclear RNA biology is not unprecedented as it was previously shown to repress nuclear mRNA export and alter nuclear RNA processing^23,25^. Thus, mass decay of cytosolic mRNAs signals via yet-to-be-determined mechanisms to repress both host gene expression via inhibition of RNAPII-mediated transcription, perturbation of nuclear RNA processing, and inhibition of nuclear mRNA export. This likely serves to reinforce the ability of RNase L to suppress the homoeostatic proteome and autoregulate inflammatory responses. It is notable that despite bulk reduction in RNAPII-mediated transcription at 8 hrs. post-lipofection of poly(I:C), ISGs such as *IFIT2* were being actively transcribed based on the presence of abundant mRNA and nascent mRNAs at the genomic locus (**Figure 6D**). These observations suggest that the repression of RNAPII-mediated transcription by RNase L is hierarchical with respect to the kinetics by which genes are transcriptionally silenced, whereby constitutively expressed genes are preferentially repressed while stress response genes maintain induction during the initial response. Based on studies of muSOX-mediated mRNA decay,^29,32,33^ we suspect that the influx of RNA-binding proteins into the nucleus interferes with RNAPII-mediated transcription and/or RNA processing.^23^ An alternate model is that the downregulation of transcription factors by RNase L is responsible for the decrease in transcription (**Figure 3F**). Future work will aim to address the mechanism of RNase L-mediated repression of RNAPII and how this impacts viral and host gene expression.

Lastly, we identified hundreds of proteins that are differentially regulated in an RNase L-dependent manner during the innate antiviral response (**Figure 3B,C**). Many proteins that were downregulated by RNase L due to their inherently short half-life (**Figure 3D**), and they function in regulating homeostatic activities such as cell division and protein synthesis (**Figure 3E**), thus providing another means by which RNase L shuts down cellular functions. In addition, RNase L regulated proteins that have context-specific antiviral activities, such as cell-mediated cytotoxic immune responses, complement system, barrier immunity, and immune checkpoints. Future studies will aim to identify the key factors regulated by RNase L that impact antiviral immunity and tumor suppression in biologically relevant contexts.

## MATERIALS & METHODS

### Cell Culture

A549 cells were cultured in DMEM +10% fetal bovine serum (FBS) + penicillin/streptomycin (P/S). Passaging used 0.25% trypsin. C6/36 and Vero cell lines were cultured in EMEM (cat #30-2003) + 10% FBS + P/S. CCL125 cells were cultured in EMEM +20% FBS + P/S. C6/36 and CCL125 cells were cultured at 28C with 5% CO2. A549, Vero, and U2OS cells were cultured at 37C with 5% CO2.

### Generation and quantification of viral stock

Viral stocks were produced in Vero cells. Once the cells were >80% confluent, cells were infected at 0.1 MOI. Supernatants were harvested at either 48 or 72 hours. After collecting the supernatant, dead cells were pelleted by centrifugation at 500g for 5 minutes. The remaining media was syringe filtered using a 0.45 uM filter.

### Plaque assays

#### Viral infections

Cells were plated one day before infection and grown to 60-80% confluency. Viruses were added in serum-free media at indicated MOIs. After one hour, media was removed, cells were washed 1x with Dulbecco’s phosphate-buffered saline (PBS), and fresh media was added.

### smFISH probe labeling

Probes were designed using Stellaris probe designer tool (Supplemental data file). DNA oligos (Integrated DNA Technologies) were labeled with 5-Propargylamino-ddUTP (ATTO-633, ATTO-550, or ATTO-488) using Terminal Deoxynucleotidyl Transferase (Thermo Fisher Scientific) according to manufacturer recommendation.

### Microscopy

Cells were plated onto glass coverslips in 12-well plates, then virally infected. After the indicated time of infection, the media was removed, cells were washed 1x in DPBS, and then fixed in 500uL 4% paraformaldehyde for 12 minutes. To permeabilize the cells, pfa was then removed, and 1mL 75% ethanol was added. Cells were then stored at 4C for at least two hours before beginning staining protocol. For dual staining of immunofluorescence and smFISH, cells were washed 2x in PBS, then incubated in 500uL primary antibody in PBS for four hours at 4C. Cells were washed 2x in PBS, then secondary antibodies were added for two hours at 4C. Cells were washed 2x, then fixed in 4% pfa for 10 minutes. Cells were washed 3x in PBS, then washed in buffer A (filter-sterilized 2x SSC with 10% formamide) for 5 minutes. smFISH probes were then added to a hybridization chamber containing a wet paper towel with parafilm on top. 50uL of smFISH probes diluted 1:100 in hybridization buffer (0.45um filtered 10% dextran sulfate, 10% formamide, 1x nuclease-free SSC, diluted in nuclease-free water) were dropped onto the parafilm. Glass slips were then flipped onto the smFISH probes. Hybridization chambers were sealed with parafilm and incubated overnight at 37C. The next day, slips were washed 2x in Buffer A, then once in 2x SSC. Slips were then mounted on slides with Vectashield + DAPI (VectorLabs: H-1000-10) and dried. Images were taken on a Nikon Eclipse Ti2 with a 100x, 1.45 NA oil objective or a 20x air objective.

### SP3 Peptide Cleanup

Frozen cell pellets were lysed with SDS lysis buffer: 50mM HEPES pH 8.5, 1% SDS, protease inhibitor, and 500 U/mL Pierce Universal Nuclease (ThermoFisher #88700). Samples were homogenized with 20 strokes of a Dounce homogenizer and then incubated at 4C for 30min. The lysate was clarified by centrifugation at 20,000rcf for 10min and the supernatant was transferred to a new tube. Samples were normalized to 1mg/ml. 20 µg of protein from each sample was prepared for LC/MS analysis using Sera-Mag Carboxylate SpeedBeads (Cytiva Life Sciences, Cat. #45152105050250 and Cat. #65152105050250) following the manufacturer protocol. In brief, samples were reduced and alkylated with 10mM DTT and 22.5mM iodoacetamide. Samples were loaded onto Sera-Mag Carboxylate SpeedBeads at 1:20 protein:bead ration. 100% Ethanol was used to bind protein to bead, followed by 80% Ethanol washes. Proteins were digested at 37C overnight with Pierce Trypsin Protease MS-Grade (Thermo Scientific #90057). Trypsinized peptides were then cleaned up with Sera-Mag Carboxylate SpeedBeads, using acetonitrile to bind and subsequently wash samples. Peptides were eluted with 2% DMSO in water and sonicated for 1 minute. Eluted peptides were split into two technical replicates per sample and submitted to The Herbert Wertheim UF Scripps Institute for Biomedical Innovation and Technology’s mass spectrometry and proteomics core and run on a Bruker timsTOF Pro2

### Mass Spectrometry Data Analysis

Peptide digests were acidified with TFA to 0.1% (v:v) and desalted using 2 μg capacity ZipTips (Millipore, Billerica, MA) according to manufacturer instructions. Following drying under vacuum, peptides were re-solubilized in 20mL of 0.1% formic acid (FA) to a final concentration of 100ng/µL. Samples were analyzed on a nanoElute (plug-in V2.1.60.0; Bruker, Germany) coupled to a Bruker TimsTOF Pro 2 mass spectrometer (Breman, Germany), equipped with a CaptiveSpray source and a 20μm zero dead volume (ZDV) Sprayer. Peptides (corresponding to 400ng) were separated on a reverse-phase C18 column (10cm X 75μm X 1.9μm, Bruker PepSep Ten Series). The column temperature was maintained at 50°C using an integrated Bruker Column Toaster (Bremen, Germany). The column was equilibrated using 4 column volumes before loading samples in 100% buffer A (99.9% Fisher Optima® LC/MS water, 0.1% FA), with both steps performed at 800 bar. Samples were separated at 500nl/min using a linear gradient from 3% to 30% buffer B (99.9% Fisher Optima® LC/MS acetonitrile, 0.1% FA) over 17.90min before ramping to 95% buffer B (0.5min) and sustained at 95% buffer B for 2.4min (total separation method time 20.7min). The Bruker TimsTOF Pro 2 was operated in DIA-PASEF mode using Tims Control v. 5.0.2. Settings for the MS method were as follows: Mass Range 100 to 1700m/z, 1/K0 Start 0.6 V.·/cm2 End 1.4 V·s/cm2, TIMS Ramp and accumulation time 75ms, Capillary Voltage 1700V, Dry Gas 3 l/min, Dry Temp 200°C, DIA-PASEF settings: 18 MS/MS scans (50m/z windons, 0.21 1/K0 windows, total cycle time 0.74), mass range 300 to 1200, and CID collision energy 20eV (at 0.60, 1/K0) to 65eV (at 1.60, 1/K0). The analysis was performed at The Herbert Wertheim UF Scripps Institute for Biomedical Innovation & Technology, Mass Spectrometry and Proteomics Core Facility (RRID:SCR_023576).

Data was processed via DIANN 1.8.1. Parameters set as follows: trypsin/P digestion, 3 missed cleavages, 3 max. variable modifications, N-term M excision, Ox(M), Ac(N-term) and C carbamidomethylation. Peptide length range was 7-30, precursor charge range 1-4, m/z range 300-1800, and fragment ion range 200-1800. Mass accuracy and MS accuracy were both set to 10. The following settings on the algorithm were checked: “Use isotopologues”, “MBR”, “No shared spectra”, “Heuristic protein inference”. Precursor FDR was set to 1%. A spectral library was used generated via DIANN from all known human proteins (In-Silico specral library). Resulting matrix.pg file was opened in Perseus (v2.0.7.0). Intensities were inputted as “main”, the rest of the descriptors are categorical. Data was then transformed (Log_2_). Data was annotated as by sample, grouping all replicates together. Missing values were imputed with Perseus default settings. Normalization was performed via median subtraction. Following this process, a volcano plot was generated utilizing a t-test for statistical significance. The resulting volcano plots were plotted in GraphPad Prism 10 for final figures. Gene Ontology was conducted using Metascape, and the data was plotted using ggplot in R.

### Statistical analyses

P-values were derived either by one-way ANOVA with Tukey HSD (https://www.socscistatistics.com/tests/anova/default2.aspx) or student’s t-test (Excel). The specific test used for each figure is specified in each figure legend. * p<0.05, **p>0.01, ***p>0.001 unless otherwise noted in the figure legend.

## ACKNOWLEDGMENTS

We thank Dr. Hyeryun Choe and Dr. Lizhou Zhang for providing the virus stocks (Boston Children’s Hospital). This work was funded by institutional funds from the Herbert Wertheim University of Florida Scripps Institute for Biomedical Innovation and Technology (JMB) and by the Office of The Director, of the National Institutes of Health under Award Number S10OD036363, and the National Institute of General Medical Sciences of the National Institutes of Health R35GM151249 (JMB), R35GM150765 (CPS). The content is solely the responsibility of the authors and does not necessarily represent the official views of the national institutes of health.

## AUTHOR CONTRIBUTIONS

JMW and CJD initiated the study. JWM performed ZIKV infections. CJD performed MS. RC performed smRNA-FISH and IFA analyses. JMW, CJD, RC, CS, and JMB analyzed data. JMW, JMB and CS wrote the manuscript.

## COMPETING INTEREST STATEMENT

The authors declare no competing interests.

## Notes

### Competing Interest Statement

The authors have declared no competing interest.

## REFERENCES

1. Bisbal, C., and Silverman, R.H. (2007). Diverse functions of RNase L and implications in pathology. Biochimie 89, 789–798. 10.1016/j.biochi.2007.02.006.

2. Hovanessian, A.G., Brown, R.E., and Kerr, I.M. (1977). Synthesis of low molecular weight inhibitor of protein synthesis with enzyme from interferon-treated cells. Nature 268, 537–540. 10.1038/268537a0.

3. Kerr, I.M., and Brown, R.E. (1978). pppA2’p5’A2’p5’A: an inhibitor of protein synthesis synthesized with an enzyme fraction from interferon-treated cells. Proc Natl Acad Sci USA 75, 256–260. 10.1073/pnas.75.1.256.

4. Clemens, M.J., and Williams, B.R. (1978). Inhibition of cell-free protein synthesis by pppA2’p5’A2’p5’A: a novel oligonucleotide synthesized by interferon-treated L cell extracts. Cell 13, 565–572. 10.1016/0092-8674(78)90329-x.

5. Baglioni, C., Minks, M.A., and Maroney, P.A. (1978). Interferon action may be mediated by activation of a nuclease by pppA2’p5’A2’p5’A. Nature 273, 684–687. 10.1038/273684a0.

6. Han, Y., Donovan, J., Rath, S., Whitney, G., Chitrakar, A., and Korennykh, A. (2014). Structure of human RNase L reveals the basis for regulated RNA decay in the IFN response. Science 343, 1244–1248. 10.1126/science.1249845.

7. Wreschner, D.H., James, T.C., Silverman, R.H., and Kerr, I.M. (1981). Ribosomal RNA cleavage, nuclease activation and 2-5A(ppp(A2’p)nA) in interferon-treated cells. Nucleic Acids Res. 9, 1571–1581. 10.1093/nar/9.7.1571.

8. Floyd-Smith, G., Slattery, E., and Lengyel, P. (1981). Interferon action: RNA cleavage pattern of a (2’-5’)oligoadenylate--dependent endonuclease. Science 212, 1030–1032. 10.1126/science.6165080.

9. Silverman, R.H. (2007). Viral encounters with 2’,5’-oligoadenylate synthetase and RNase L during the interferon antiviral response. J. Virol. 81, 12720–12729. 10.1128/JVI.01471-07.

10. Chakrabarti, A., Jha, B.K., and Silverman, R.H. (2011). New insights into the role of RNase L in innate immunity. J. Interferon Cytokine Res. 31, 49–57. 10.1089/jir.2010.0120.

11. Min, J.-Y., and Krug, R.M. (2006). The primary function of RNA binding by the influenza A virus NS1 protein in infected cells: Inhibiting the 2’-5’ oligo (A) synthetase/RNase L pathway. Proc Natl Acad Sci USA 103, 7100–7105. 10.1073/pnas.0602184103.

12. Zhao, L., Jha, B.K., Wu, A., Elliott, R., Ziebuhr, J., Gorbalenya, A.E., Silverman, R.H., and Weiss, S.R. (2012). Antagonism of the interferon-induced OAS-RNase L pathway by murine coronavirus ns2 protein is required for virus replication and liver pathology. Cell Host Microbe 11, 607–616. 10.1016/j.chom.2012.04.011.

13. Li, Y., Banerjee, S., Wang, Y., Goldstein, S.A., Dong, B., Gaughan, C., Silverman, R.H., and Weiss, S.R. (2016). Activation of RNase L is dependent on OAS3 expression during infection with diverse human viruses. Proc Natl Acad Sci USA 113, 2241–2246. 10.1073/pnas.1519657113.

14. Thornbrough, J.M., Jha, B.K., Yount, B., Goldstein, S.A., Li, Y., Elliott, R., Sims, A.C., Baric, R.S., Silverman, R.H., and Weiss, S.R. (2016). Middle east respiratory syndrome coronavirus ns4b protein inhibits host rnase L activation. MBio 7, e00258. 10.1128/mBio.00258-16.

15. Malathi, K., Paranjape, J.M., Ganapathi, R., and Silverman, R.H. (2004). HPC1/RNASEL mediates apoptosis of prostate cancer cells treated with 2’,5’-oligoadenylates, topoisomerase I inhibitors, and tumor necrosis factor-related apoptosis-inducing ligand. Cancer Res. 64, 9144–9151. 10.1158/0008-5472.CAN-04-2226.

16. Malathi, K., Dong, B., Gale, M., and Silverman, R.H. (2007). Small self-RNA generated by RNase L amplifies antiviral innate immunity. Nature 448, 816–819. 10.1038/nature06042.

17. Chakrabarti, A., Ghosh, P.K., Banerjee, S., Gaughan, C., and Silverman, R.H. (2012). RNase L triggers autophagy in response to viral infections. J. Virol. 86, 11311–11321. 10.1128/JVI.00270-12.

18. Chakrabarti, A., Banerjee, S., Franchi, L., Loo, Y.-M., Gale, M., Núñez, G., and Silverman, R.H. (2015). RNase L activates the NLRP3 inflammasome during viral infections. Cell Host Microbe 17, 466–477. 10.1016/j.chom.2015.02.010.

19. Burke, J.M., Moon, S.L., Matheny, T., and Parker, R. (2019). RNase L Reprograms Translation by Widespread mRNA Turnover Escaped by Antiviral mRNAs. Mol. Cell 75, 1203–1217.e5. 10.1016/j.molcel.2019.07.029.

20. Rath, S., Prangley, E., Donovan, J., Demarest, K., Wingreen, N.S., Meir, Y., and Korennykh, A. (2019). Concerted 2-5A-Mediated mRNA Decay and Transcription Reprogram Protein Synthesis in the dsRNA Response. Mol. Cell 75, 1218–1228.e6. 10.1016/j.molcel.2019.07.027.

21. Cusic, R., and Burke, J.M. (2024). Condensation of RNase L promotes its rapid activation in response to viral infection in mammalian cells. Sci. Signal. 17, eadi9844. 10.1126/scisignal.adi9844.

22. Chitrakar, A., Rath, S., Donovan, J., Demarest, K., Li, Y., Sridhar, R.R., Weiss, S.R., Kotenko, S.V., Wingreen, N.S., and Korennykh, A. (2019). Real-time 2-5A kinetics suggest that interferons β and λ evade global arrest of translation by RNase L. Proc Natl Acad Sci USA 116, 2103–2111. 10.1073/pnas.1818363116.

23. Burke, J.M., Gilchrist, A.R., Sawyer, S.L., and Parker, R. (2021). RNase L limits host and viral protein synthesis via inhibition of mRNA export. Sci. Adv. 7. 10.1126/sciadv.abh2479.

24. Karasik, A., Lorenzi, H.A., DePass, A.V., and Guydosh, N.R. (2024). Endonucleolytic RNA cleavage drives changes in gene expression during the innate immune response. Cell Rep. 43, 114287. 10.1016/j.celrep.2024.114287.

25. Burke, J.M., Ripin, N., Ferretti, M.B., St Clair, L.A., Worden-Sapper, E.R., Salgado, F., Sawyer, S.L., Perera, R., Lynch, K.W., and Parker, R. (2022). RNase L activation in the cytoplasm induces aberrant processing of mRNAs in the nucleus. PLoS Pathog. 18, e1010930. 10.1371/journal.ppat.1010930.

26. Zhou, A., Paranjape, J., Brown, T.L., Nie, H., Naik, S., Dong, B., Chang, A., Trapp, B., Fairchild, R., Colmenares, C., et al. (1997). Interferon action and apoptosis are defective in mice devoid of 2’,5’-oligoadenylate-dependent RNase L. EMBO J. 16, 6355–6363. 10.1093/emboj/16.21.6355.

27. Xi, J., Snieckute, G., Martínez, J.F., Arendrup, F.S.W., Asthana, A., Gaughan, C., Lund, A.H., Bekker-Jensen, S., and Silverman, R.H. (2024). Initiation of a ZAKα-dependent ribotoxic stress response by the innate immunity endoribonuclease RNase L. Cell Rep. 43, 113998. 10.1016/j.celrep.2024.113998.

28. Burke, J.M., Lester, E.T., Tauber, D., and Parker, R. (2020). RNase L promotes the formation of unique ribonucleoprotein granules distinct from stress granules. J. Biol. Chem. 295, 1426–1438. 10.1074/jbc.RA119.011638.

29. Gilbertson, S., Federspiel, J.D., Hartenian, E., Cristea, I.M., and Glaunsinger, B. (2018). Changes in mRNA abundance drive shuttling of RNA binding proteins, linking cytoplasmic RNA degradation to transcription. eLife 7. 10.7554/eLife.37663.

30. Decker, C.J., Burke, J.M., Mulvaney, P.K., and Parker, R. (2022). RNA is required for the integrity of multiple nuclear and cytoplasmic membrane-less RNP granules. EMBO J. 41, e110137. 10.15252/embj.2021110137.

31. Banerjee, S., Chakrabarti, A., Jha, B.K., Weiss, S.R., and Silverman, R.H. (2014). Cell-type-specific effects of RNase L on viral induction of beta interferon. MBio 5, e00856–14. 10.1128/mBio.00856-14.

32. Kumar, G.R., and Glaunsinger, B.A. (2010). Nuclear import of cytoplasmic poly(A) binding protein restricts gene expression via hyperadenylation and nuclear retention of mRNA. Mol. Cell. Biol. 30, 4996–5008. 10.1128/MCB.00600-10.

33. Abernathy, E., Gilbertson, S., Alla, R., and Glaunsinger, B. (2015). Viral Nucleases Induce an mRNA Degradation-Transcription Feedback Loop in Mammalian Cells. Cell Host Microbe 18, 243–253. 10.1016/j.chom.2015.06.019.

