## Supplemental Figures for "RNase L regulates the antiviral proteome by accelerating mRNA decay, inhibiting nuclear mRNA export, and repressing RNAPII-mediated transcription"

### Supplemental figure legends

A

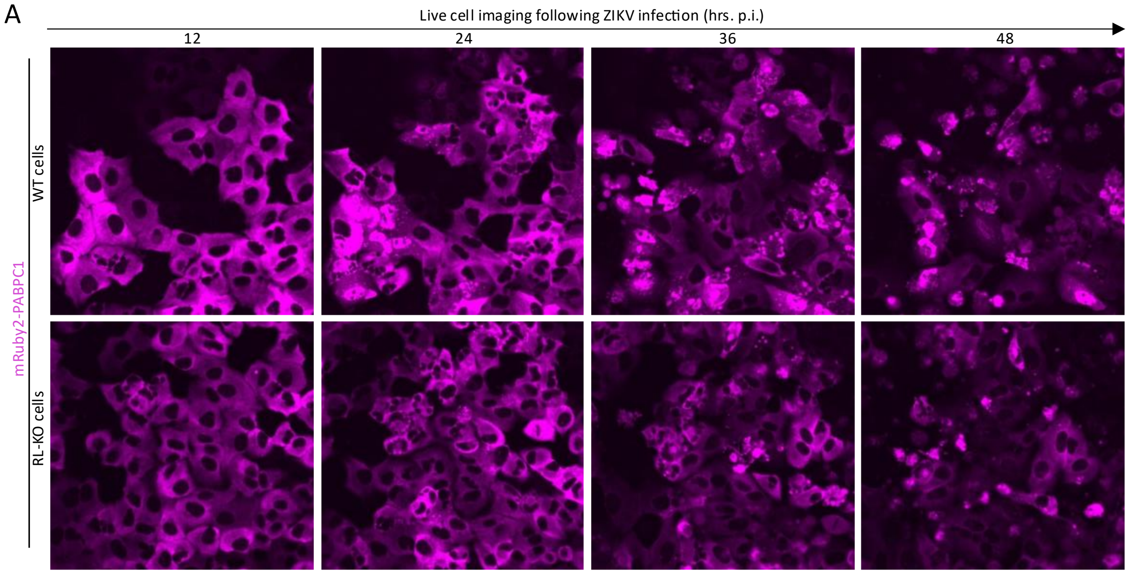


**Figure S1 (related to Fig. 2). RNase L-dependent regulation of the cellular proteome during dsRNA stress**

(A) Livecell imaging of WT and RLKO mRUBY1-PABPC1 cells during ZIKV infection.

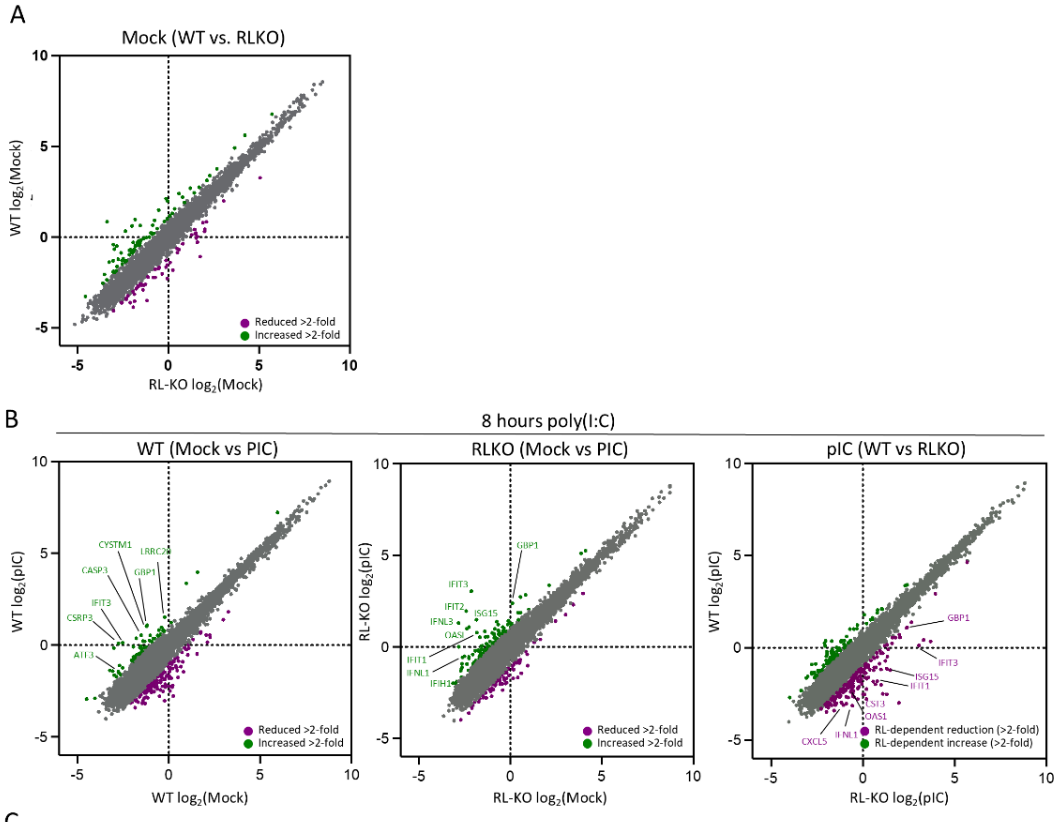


**Figure S2 (related to Fig. 3). RNase L-dependent regulation of the cellular proteome during dsRNA stress**

(A) Scatter plot of quantitative MS in WT and RL-KO cells under mock conditions. (B) Scatter plot analyses of protein levels in mock conditions (x-axis) and 8 hours post-lipofection of poly(I:C) in WT or RL-KO cells.

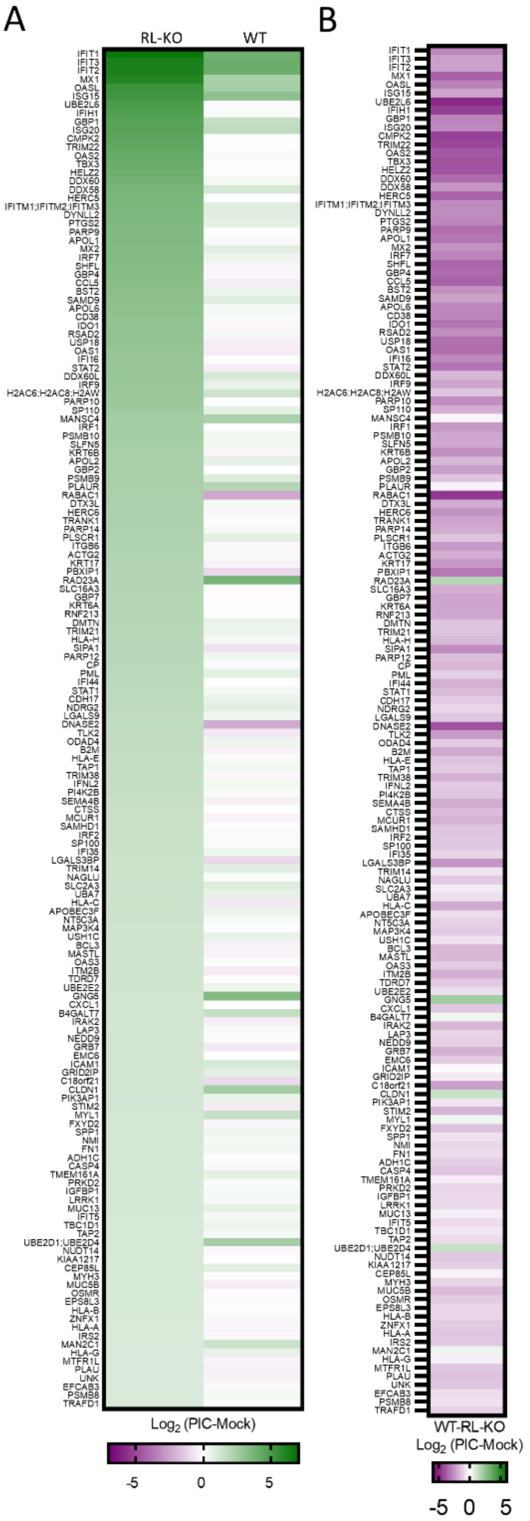


**Figure S3 (related to Fig. 3). RNase L-dependent alterations to dsRNA-induced proteins.** (A) Heat map of proteins that are upregulated at least twofold in RL-KO cells in response to poly(I:C). (B) Heat map showing RNase L-dependent changes to proteins in (A).

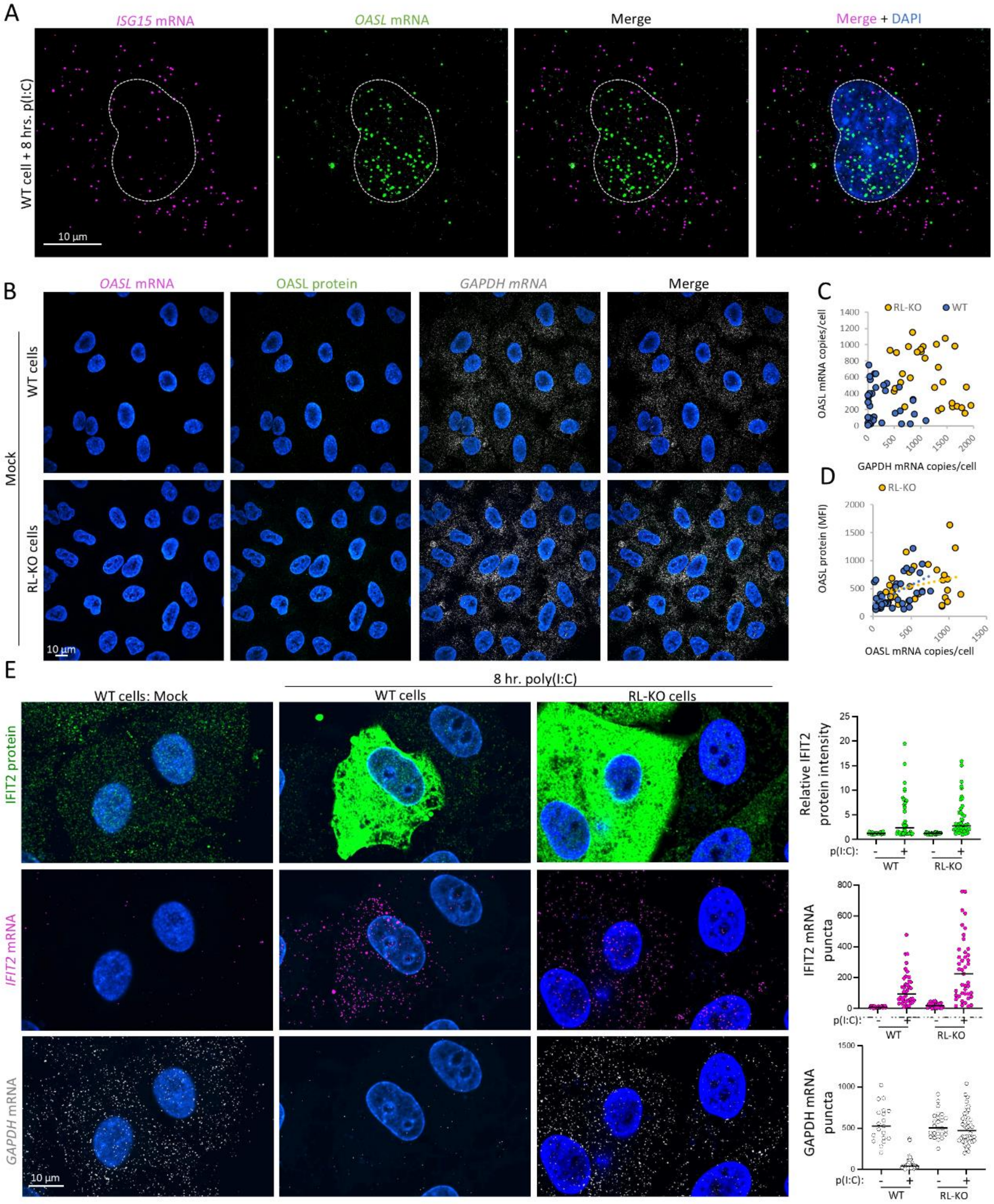


**Figure S4 (related to Fig. 4). Antiviral protein synthesis during the RNase L response.** (A) smRNA-FISH for *ISG15* and *OASL* mRNAs. (B) IFA for OASL protein and smRNA-FISH for *GAPDH* and *OASL* mRNAs under mock conditions in WT and RL-KO cells. (C) Scatter plot analysis of OASL mRNA copies (y-axis) and *GAPDH* mRNA copies (x-axis) in WT and RL-KO cells. (D) Scatter plot analysis of OASL protein mean fluorescence intensity (y-axis) and *OASL* mRNA copies (x-axis) in WT and RL-KO cells. (E) IFA for IFIT2 protein and smRNA-FISH for *IFIT2* and *OASL* mRNAs under mock conditions in WT and RL-KO cells. The graphs on the right display the quantification from individual cells as represented in the image panels.

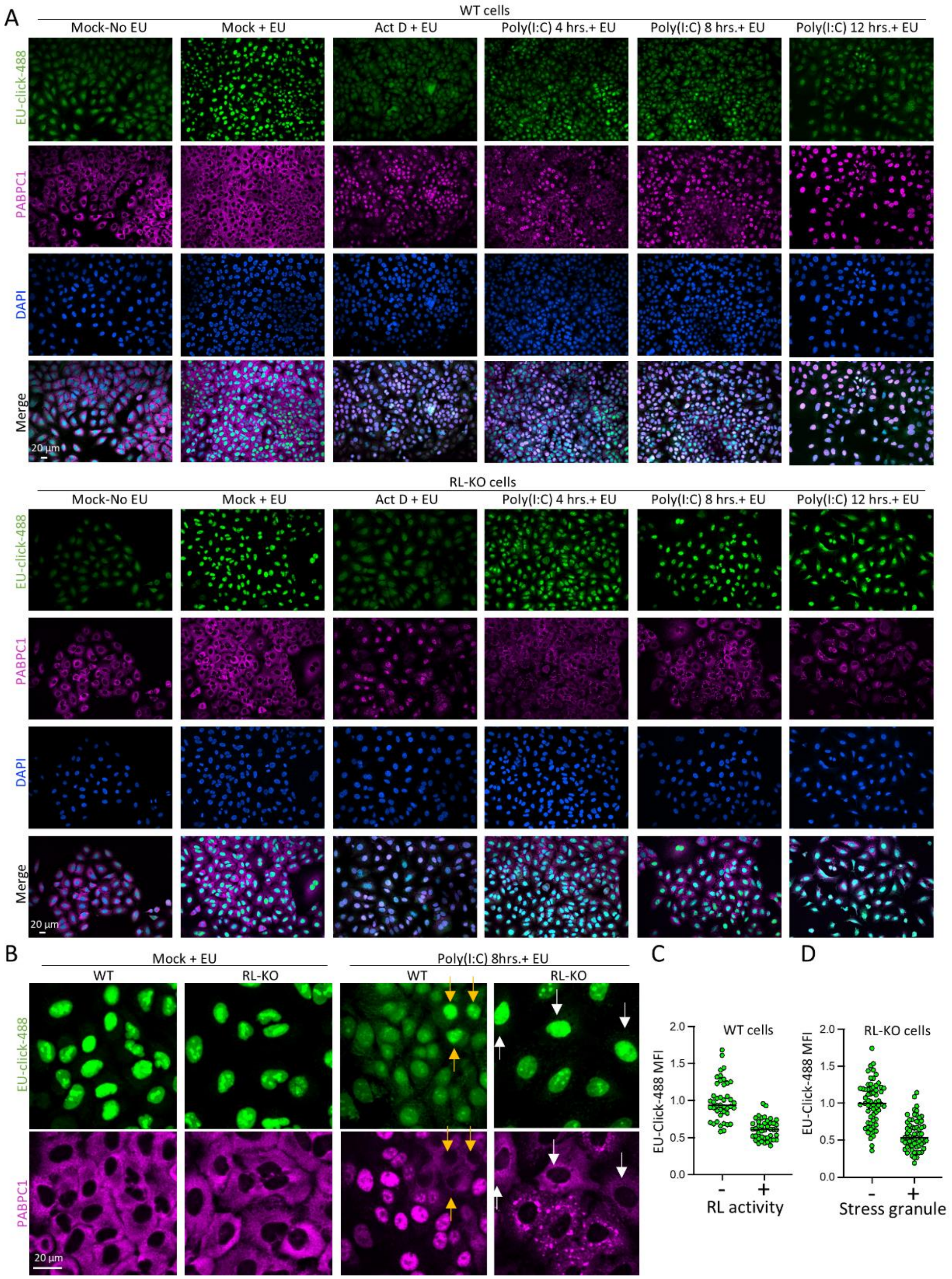


**Figure S5 (related to Fig. 6). EU metabolic labeling of RNA.**
(A) Microscopy of EU-click-488 fluorescence and IFA for PABPC1 in WT and RL-KO cells. (B) Inset from panels in (A) showing PAPBC1 localization. (C) Normalized EU mean fluorescence intensity (MFI) in WT cells with activate RNase L (PABPC1 in the nucleus) or inactive RNase L (PABPC1 in the cytosol). (D) Normalized EU mean fluorescence intensity (MFI) in RL-KO cells with or without PABPC1-positivie stress granules.

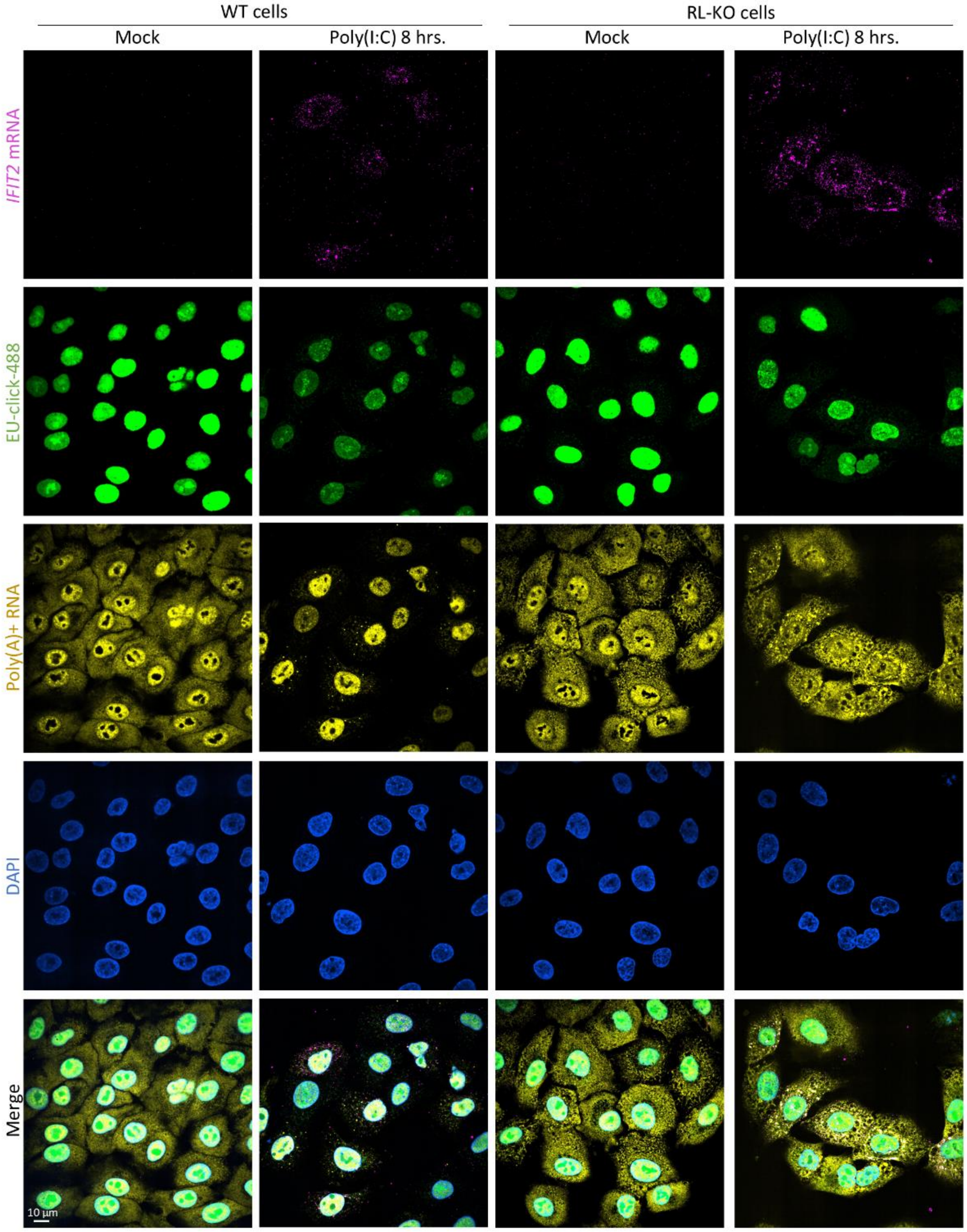


**Figure S6 (related to Fig. 6). EU metabolic labeling of RNA and smRNA-FISH for IFIT2 mRNA.**
smRNA-FISH for IFIT2 mRNA and poly(A) + RNA in WT or RL-KO cells labeled with EU-click-488 8 hours post-lipofection of poly(I:C).

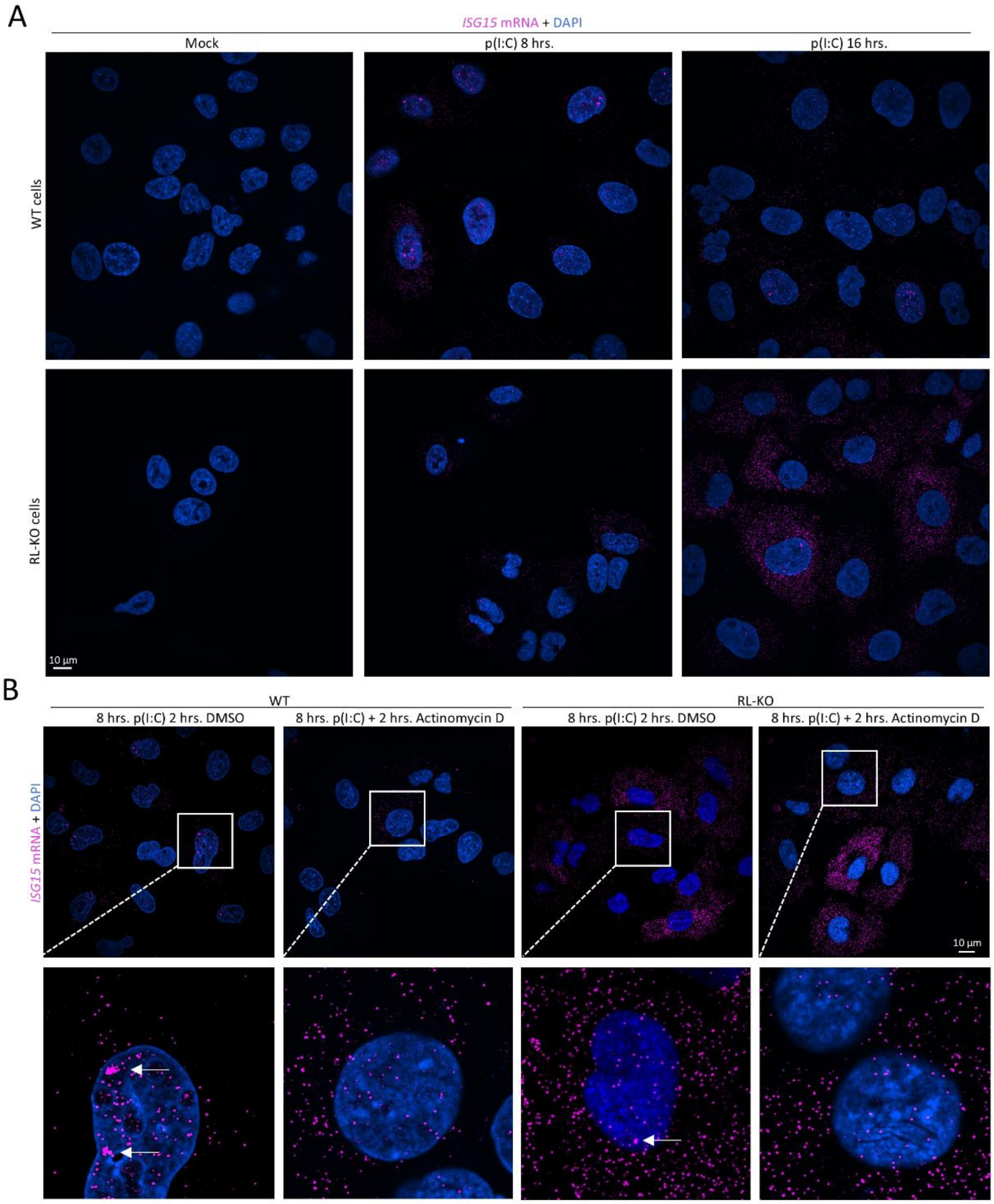


**Figure S7 (Related to figure 6). Inhibition of transcription abolishes ISG15 transcription foci.**

(A) smRNA-FISH for ISG15 mRNA in WT and RL-KO cells under mock conditions or 8- or 16-hrs. post-lipofection with poly(I:C). (B) smRNA-FISH for ISG15 mRNA in WT and RL-KO cells lipofected with poly(I:C) for 8 hours. Cells were then treated with or without (DMSO-only) actinomycin D for 2 hours. Arrows indicate *ISG15* transcription foci.
